# Influenza A Virus Coinfection Alters *Streptococcus pneumoniae* Gene Expression during Upper Respiratory Tract Colonization

**DOI:** 10.64898/2026.05.19.726214

**Authors:** Noah A. Nutter, Alicia Costa-Terryll, Lance M. Miller, Ming Leung, Pengbo Zhang, Daniel Fecko, M. Ammar Zafar

## Abstract

*Streptococcus pneumoniae* (*Spn*) asymptomatically colonizes the upper respiratory tract (URT), a niche from which it can transmit to another host or cause invasive disease in the same host. The *in vivo* transcriptional adaptations that *Spn* undergoes during nasopharyngeal colonization, particularly during influenza A virus (IAV) coinfection, are poorly understood. Here, we leveraged an established infant mouse model of colonization, shedding, and transmission to perform genome-wide transcriptomic profiling of *Spn* during mono- and during IAV coinfection. Compared with broth-grown controls, pneumococci isolated from the URT exhibited distinct transcriptional programs, with over 200 genes differentially expressed across time points. Genes involved in carbohydrate uptake and metabolism, glycan degradation, amino sugar and nucleotide sugar metabolism, and amino acid biosynthesis were consistently enriched during colonization, highlighting metabolic adaptation to the nasopharyngeal niche. In contrast, IAV coinfection induced a markedly distinct transcriptional signature, including upregulation of branched-chain amino acid biosynthesis, bacteriocin production, and phosphate acquisition systems. Notably, the pilus islet-1 locus was upregulated during *Spn*-IAV coinfection. Functional studies demonstrated that while the pilus was dispensable for colonization under mono- and coinfection conditions, it promoted high-shedding events and enhanced inflammatory responses during IAV coinfection. However, reduced inflammation and reduced high shedding events from pups inoculated with a pilus-deficient mutant did not alter transmission frequency in the infant mouse model. Collectively, our findings define the *in vivo* transcriptional landscape of *Spn* during URT colonization and reveal distinct bacterial adaptations during viral coinfection, providing insight into mechanisms that influence pneumococcal persistence, inflammation, and transmission.

## Introduction

*Streptococcus pneumoniae* (the pneumococcus; *Spn*) is the leading cause of community acquired pneumonia that mainly affects the young, elderly and the immunocompromised. The acquisition of pneumococcus in the upper respiratory tract and persistence is generally considered an asymptomatic event, and an individual can have multiple *Spn* colonizing events through their lifespan. However, from this nasopharyngeal niche *Spn* can at times gain access to sterile sites within the host and cause invasive disease including otitis media, pneumonia, sepsis and meningitis (1). The advent of the conjugate capsular polysaccharide vaccine not only led to the reduction in the number of invasive disease cases but also reduced the transmission rates between individuals, thus providing herd immunity (2, 3). While the conjugate vaccine has been highly successful, it only covers the most common disease-causing isolates, with isolates also serotype (capsule) switching to bypass the vaccine-based immunity (4, 5). Collectively, this leads to a high pneumococcal disease burden (6), requiring further study of this highly adapted pathogen.

Humans are the primary reservoir for *Spn*, with its genome highly adapted to carriage (1). While the *Spn* genome is relatively small, it allows the pathobiont to rapidly adapt to the changing environment of the host, with many bacterial factors playing multiple roles (moonlighting proteins) (7), such as the neuraminidases that provide not only sialic acid as a carbon source but also help in inactivating the host complement system and promote biofilm formation (7-9). Choline binding proteins, present on the surface of *Spn* are known to modulate host innate immune response, and are also involved in biofilm formation (10) identifying biofilm formation as a critical pathogenic feature of *Spn*; pneumolysin, a key *Spn* virulence determinant factors, is the sole cytolysin and has been demonstrated to positively affect host inflammatory response, ability to form biofilm and inhibit respiratory ciliary action (11-13). Besides proteins that are multifunctional, *Spn* also utilizes multiple two-component systems to rapidly respond to the changing host environment in the URT and during invasive disease (14). One such system, YesMN was recently demonstrated to be important for zinc acquisition, N-glycoconjugate uptake and metabolism, which consequently, contributed to *Spn* host-to-host transmission(15).

In its nasopharyngeal niche *Spn*, encounters the host microbiota and other invading pathogens (16-18), and epidemiological studies have demonstrated that *Spn* coinfections with influenza A virus (IAV) are a common occurrence in a community setting, with coinfections leading to increase in disease severity (19). Molecular studies illustrate that IAV infection leads to increased inflammation and mucus production, and *Spn* thrives in this altered environment (20, 21). However, recent data suggest a complex relationship, where *Spn*-IAV interactions can be antagonistic (22, 23). To that end, YesMN was observed to be critical for colonization during IAV coinfection but dispensable during *Spn* monoinfection, suggesting that the inflamed URT environment during IAV coinfection renders YesMN regulated determinants indispensable for *Spn* colonization (15).

Despite the observations described above, it remains unknown what mechanisms *Spn* employes to adapt to the nasopharynx environment in the presence or absence of IAV. Therefore, to identify the adaptive changes that *Spn* undergoes in the nasopharynx, we utilized our infant murine model of infection that recapitulates many of key elements for successful colonization and transmission of *Spn* (24), and performed a transcriptomics analysis at multiple time points post-inoculation, with and without IAV coinfection. Pathways involving glycoconjugate cleavage and uptake were highly enriched *in vivo*, with distinct transcriptional profiles observed during *Spn*-IAV coinfection that included the *Spn* type 1 pilus locus. We show that the pilus impacts inflammation and affected pneumococcal shedding without altering URT colonization. These data provide insight into the global transcriptional modulations that *Spn* undergoes during URT colonization and how IAV coinfection influences the post-inoculation transcriptome.

## Results

### Optimization of the infant mouse model for *in vivo* transcriptional analysis

Studies have indicated both a synergistic and an antagonistic relationship between *Spn* and IAV in the upper respiratory tract (URT), (15, 22, 23) suggesting that *Spn* possibly undergoes transcriptional changes during IAV infection of the URT. To determine the peak IAV associated host response during *Spn* URT colonization we performed quantitative reverse-transcription polymerase chain reaction (qRT-PCR) analysis on nasal wash samples obtained from mono and coinfected mice at different time points post IAV infection (**Fig 1A**). No major shift in *Spn* colonization density was observed while sampling the URT at different time points with and without IAV infection (**Fig S1A**). IAV infection of the URT is associated with a robust inflammatory response and increase in mucus secretions(21, 25). We assessed inflammatory transcripts CXCL-1, 2, IL1β, IFN-α and mucus (MUC5) and observed the peak inflammatory response at 72 hours post IAV inoculation (8 days post *Spn* inoculation) (**Fig S1B-D**). Thus, this time point was selected to determine the *Spn* transcriptional modulations during IAV coinfection.

**Figure 1.**
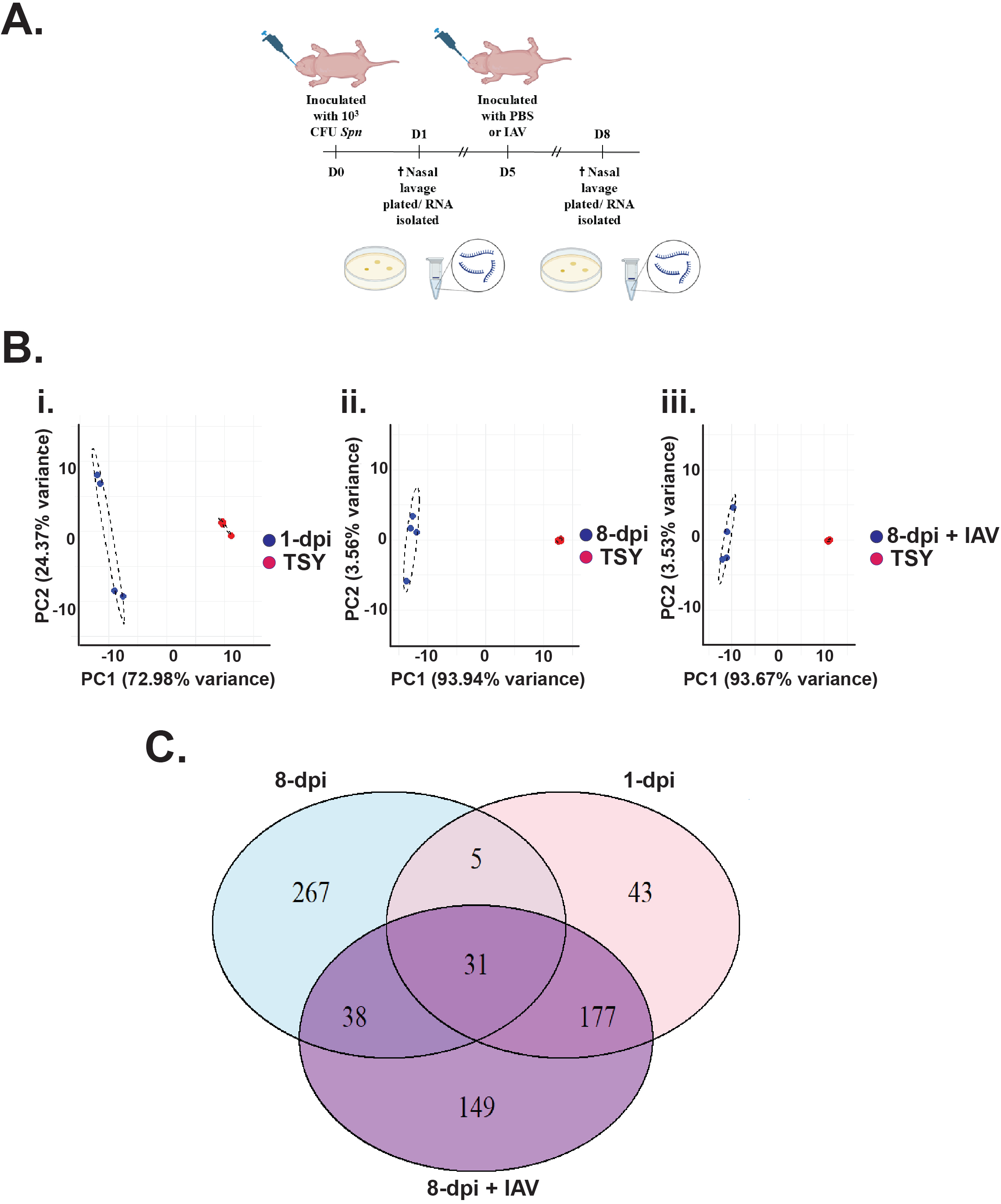
An *in vivo* RNA-seq approach provides insight into pneumococcal transcriptional changes that occur during upper respiratory tract colonization. (**A)** Schematic representation of the inoculation and nasal lavage schedule with and without a corresponding influenza A infection (IAV). Four-day-old pups (n=8 per group) were intranasally (i.n) inoculated with ∼2,000 CFU of TIGR4Sm^R^. The cross indicates the euthanization, which was performed on the designated days and nasal lavages collected and plated for determining colonization density and isolation of RNA. For *S. pneumoniae*-IAV coinfection studies, pups were initially inoculated with *S. pneumoniae* and five days post-inoculation (P.I) were either mock treated (PBS) or given IAV (2 × 10^4^ pfu) and then followed for the designated time before euthanization and nasal lavage collection. **(B)** PCA plots depicting the variance across the pneumococcal genes when compared between **(i.)** day 1 P.I and TSY, **(ii.)** day 8 P.I and TSY, and **(iii.)** day 8 P.I with a concurrent IAV infection and TSY. **(D)** Venn diagram displaying the comparison of *in vivo* differentially expressed gene expression (Log_2_FC > ±1; *P*_*adj*_ *< 0*.*05*) in comparison to TSY across different timepoints, organized into unique and overlapping expressions.

For the transcriptome analysis, *in vivo Spn* samples obtained from the URT were compared to *Spn* samples grown in tryptic soy yeast (TSY) broth. Different time points post *Spn* inoculation and *Spn*-IAV coinfection were selected to provide broad overview of the genome wide transcriptional modulations that *Spn* undergoes in the URT. The raw reads were trimmed and aligned to the pneumococcal reference genome (TIGR4)(26). Previous studies have suggested that lower read counts that quantify gene expression by mapping the sequenced RNA fragments to the reference genome are still sufficient to identify differential gene expression in animal models of bacterial infections(27-29). To circumvent the lower read counts bottleneck, total RNA was isolated from two retrograde lavage samples that were pooled together. Higher *Spn* read counts were obtained from the broth grown samples than *in vivo* samples (**Fig S2**). The lowest read counts were obtained from samples taken 24 hours post inoculation. *Spn* colonization density in the URT does not vary significantly between mice at different time points or during coinfection (**Fig S1A**) (30, 31), so the low read counts is likely due to difficulty in obtaining samples from 5-day old pups. Principal component analyses (PCA) of the top 500 differentially expressed genes (DEG) (**Table S1**) were used to determine *Spn* gene expression profiles across different time points and conditions. A distinct gene expression profile was observed when nasopharyngeal samples were compared to broth samples, where the *in vitro* samples clustered tightly together, and some variability was observed with the *in vivo* samples (**Fig 1B**). However, the nasopharyngeal samples clearly separated from the broth samples, suggesting that *Spn* transcriptional profile is different when comparing the nasopharynx versus *in vitro* growth. The number of DEG that were upregulated (log2FC≥ 1 and *padj* ≤0.05) varied between different timepoints, ranging from 256 to 341, with many of the genes being common between the three time points, with 1 day post inoculation having 43 unique genes, 8 days P.I 267 unique genes and 8 days P.I with a concurrent infection having 149 genes that were unique to that particular condition and time point. Additionally, 31 DEG’s were identified to be common across the time points tested (**Fig 1C**). Collectively, these results indicate that pneumococcus undergoes a distinct transcriptional programming during colonization of the URT.

### *Spn* regulates unique subset of genes in the upper respiratory tract

To provide further context for our results, we generated volcano plots showing all differentially expressed genes (**Fig 2A**) and heatmaps of the top 30 upregulated genes (log2 fold change ≥1) for early, late, and coinfections (**Fig 2B**) (**Table S2**). This early colonization list included genes involved in glycoconjugate cleavage and PTS sugar uptake (*SP_0060-64*; *SP_009-92*: *SP_0321-0325*) systems, glycerol uptake (*SP_2183-2185*) genes that form part of the Entner-Doudoroff pathway (*SP_0317-320*), which are involved in ketogluconate metabolism (32). Besides carbohydrate uptake and metabolism pathways, the lanthionine-containing pathway (*SP_1949-1949*) which leads to the production of peptides that are involved in bactericidal activity was also upregulated (33). The top 30 DEG from Day 1 and Day 8 P.I are highly consistent, suggesting that *Spn* generally encounters similar conditions in the nasopharynx overtime after the initial colonization event. Besides the pathways mentioned above, genes in the *ula* pathway (*SP_2031-2038*) were highly DE on Day 8 P.I. The *ula* operon is required for the uptake and metabolism of ascorbic acid, and *Spn* can utilize ascorbate as a sole carbon source (28). Several of the DEG from all three *in vivo* conditions were previously identified through transposon mutagenesis screens to be important for pneumococcal colonization (34) and transmission (30) highlighting their importance in *Spn* pathogenesis. Interestingly, the top 30 upregulated genes during *Spn*-IAV coinfection were highly dissimilar from mono-infected samples, suggesting that IAV mediated inflammation of the URT leads to further transcriptional changes in *Spn*. This list included genes involved in branch-chain amino acid (BCAA) biosynthesis (*SP_0446*), the *blp* bacteriocin (antimicrobial peptide) locus (*SP_0541*; *SP_0545*) important for bacterial competition (35), the two-component system PnpRS (*SP_2083*) involved in phosphate uptake (14), multiple ABC transporters and the serine protease HtrA (*SP_2239*), an enzyme that is highly conserved and plays a critical virulence role in streptococcal species (36-38).

**Fig 2.**
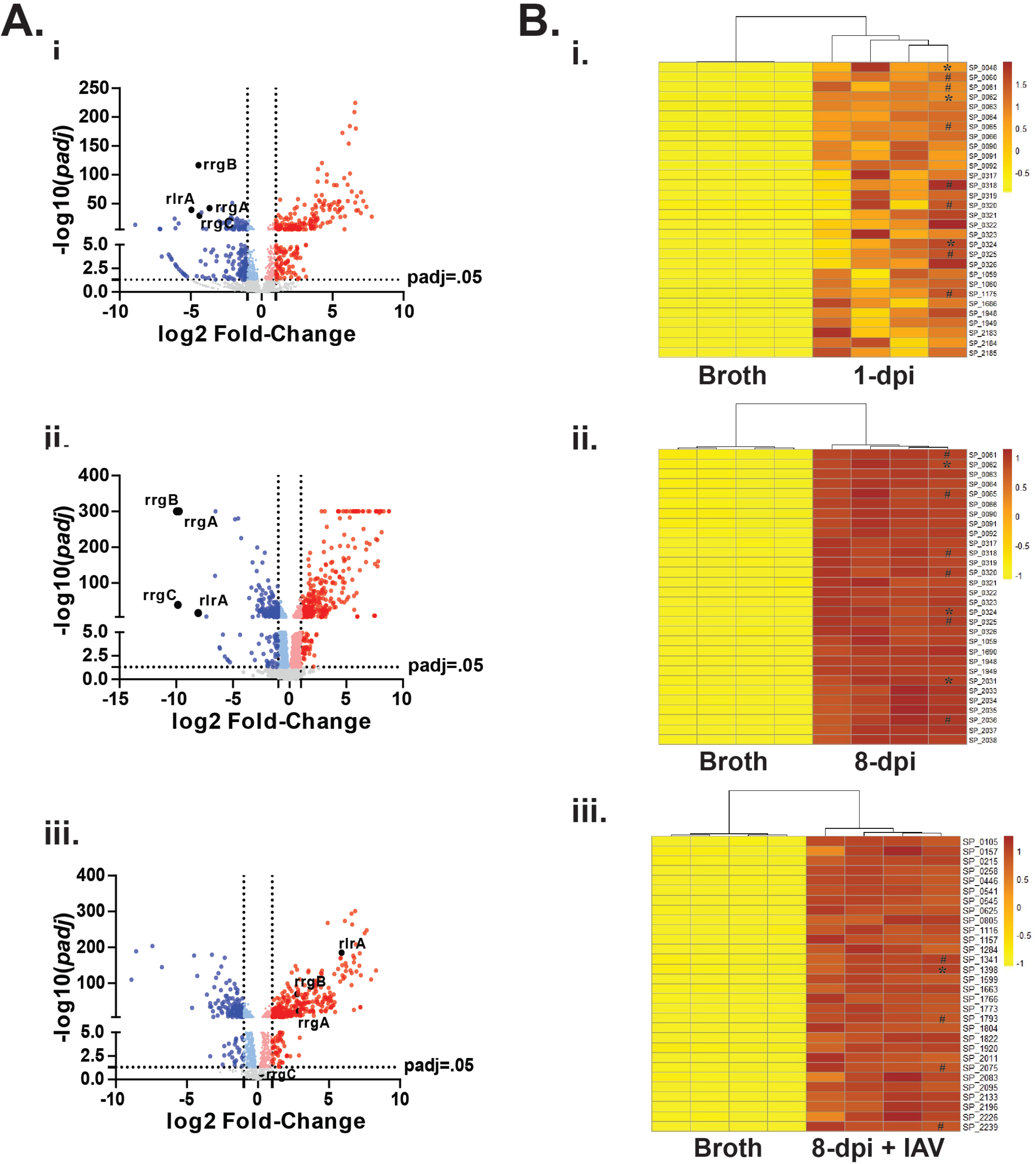
Identification of differentially regulated pneumococcal genes in upper respiratory tract. **(A)** Volcano plots depicting pneumococcal genes whose RNA transcripts were differentially regulated (Log_2_FC > ±1; *P*_*adj*_ *< 0*.*05*), **(i.)** day 1 P.I and TSY, **(ii.)** day 8 P.I and TSY, and **(iii.)** day 8 P.I with a concurrent IAV infection and TSY. where blue dots represent downregulated and red dots represent upregulated genes. Genes involved in the pilus structure are highlighted. **(B)** Heatmaps depicting the top 30 pneumococcal genes with locus tags included that are upregulated when comparing **(i.)** day 1 P.I and TSY, **(ii.)** day 8 P.I and TSY, and **(iii.)** day 8 P.I with a concurrent IAV infection and TSY. **\***represents genes previously identified to be important for pneumococcal shedding; **#**represents genes previously identified to be important for pneumococcal colonization.

Subsequently, we focused on the downregulated genes, as the 30 most downregulated genes included several loci common between the three conditions (**Fig S3**) (**Table S2**). Particularly, the pathways for cellobiose uptake and utilization (*SP_0308-SP_0309*) and trehalose uptake and utilization (*SP_1883-SP_1884*), both regulated by the global carbon catabolite control protein CcpA, were downregulated during early and late infection. Furthermore, the pilus locus (type 1 pilus) (*SP_0461-SP_0468*) present in a subset of *Spn* isolates and considered a major virulence determinant during invasive disease (39), was downregulated in both early and late infection. Thus, our results suggest that these pathways are dispensable or possibly inhibitory during colonization of the URT by *Spn*.

### Functional annotation of differentially expressed pneumococcal genes

Subsequently, to identify *Spn* molecular pathways that are required for colonization we performed gene ontology term enrichment analysis (KEGG pathway analysis) (**Fig. 3A-C**). Our enrichment analysis showed DEG’s that are involved in multiple cellular pathways including carbohydrate uptake and metabolism, amino sugar and nucleotide sugar metabolism, biosynthesis of secondary metabolites, glycan degradation, amino acid biosynthesis among others. Many of the genes belonging to an individual pathway also formed a functional component in another cellular pathway (**Fig S4)**. Several pathways that were shared across different time points include galactose metabolism, other glycan degradation, amino sugar and nucleotide metabolism, purine metabolism, biosynthesis of secondary metabolites and aminoacyl-tRNA-biosynthesis. Furthermore, many of the carbohydrate metabolism pathways altered during URT infection match those altered during *Spn* lung infection (40) or during *in vitro* alveolar epithelial cell infection (41), suggesting their overall importance to pneumococcal survival within the host. Several pathways such as thiamine metabolism, pyrimidine metabolism, pentose and glucuronate interconversions, phenylalanine, tyrosine and tryptophan biosynthesis, were only enriched in the coinfection samples. This suggests that IAV induced inflammation modulates the URT environment and pneumococcus further responds accordingly (**Fig 3C**). The aromatic amino acid biosynthesis pathway was also enriched in *Spn* associated lower respiratory tract infection (40), identifying it as a pathway important for both nasopharyngeal colonization and lung infection.

**Fig 3.**
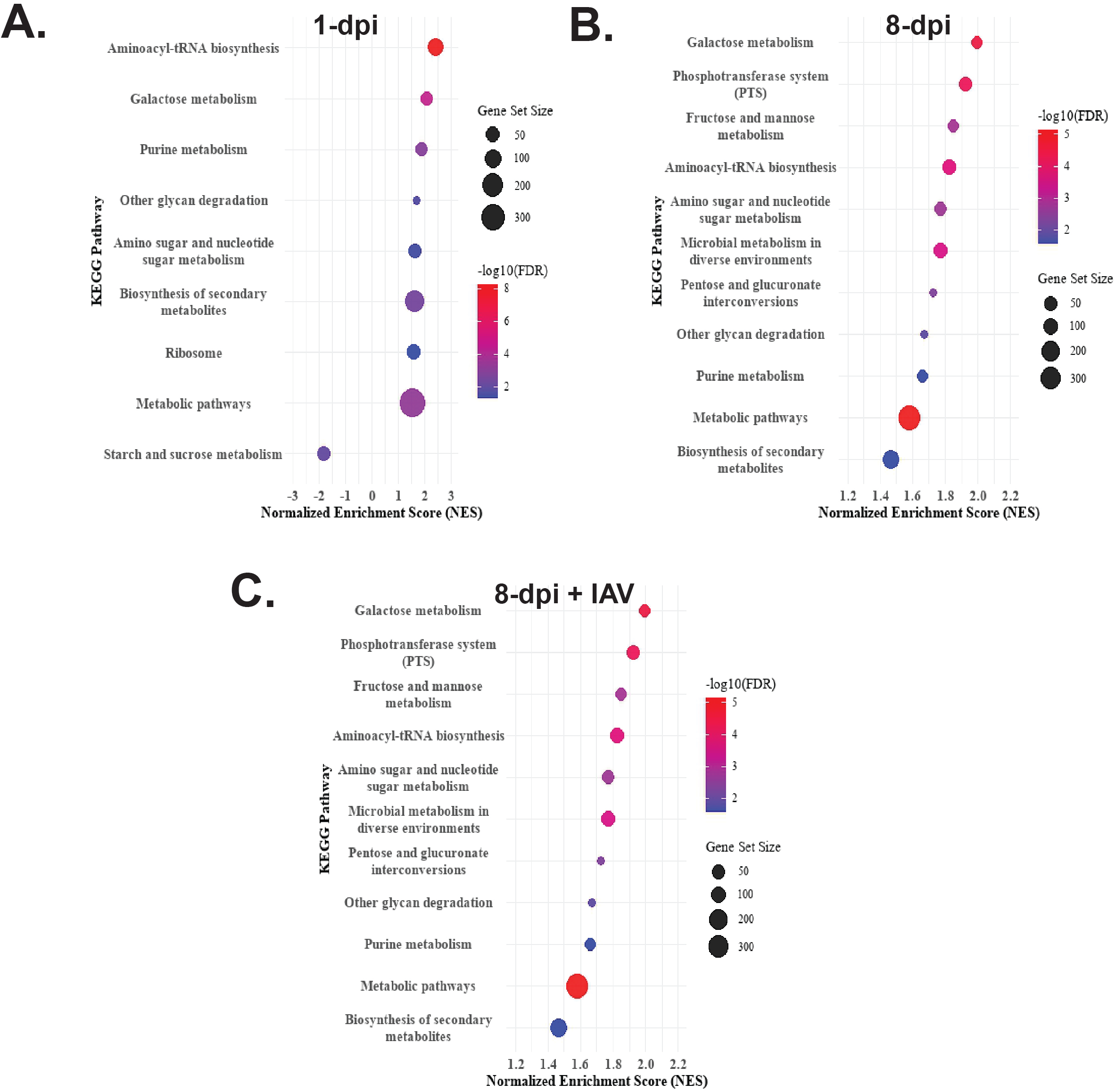
Functional annotation of the differentially regulated *S. pneumoniae* genes in the murine upper respiratory tract at different time points compared to broth. Kyoto Encyclopedia of Genes and Genomes (KEGG) pathway analyses was performed comparing **(A)** day 1 P.I to broth, **(B)** day 8 P.I to broth, and **(C)** day 8 P.I with a concurrent IAV infection (pups were given IAV at day 5 P.I to *S. pneumoniae*) to broth. A normalized enrichments score (NES) was used to determine whether a gene set is significantly enriched compared to all the genes that were upregulated. The color of the symbol is based on the adjusted significance (*Padj*) by using the false discovery rate (FDP) statistical measure with the size of the symbol (circle) representing the number of genes represented in a particular pathway.

### The *Spn* pilus locus contributes to high shedding during *Spn*-IAV coinfection

As there were considerable differences in the top 30 upregulated genes during mono- or co-infection compared to broth, we determined the differentially expressed genes between the two *in vivo* conditions (**Fig. 4A**). Included in the coinfection upregulated list was the locus (*SP_0321-0327*) involved in uptake and degradation of hyaluronate disaccharide, a key component of extracellular matrix present on the surface of epithelial cells (42). Other upregulated pathways include BCAA uptake (SP_0626; *brnQ*) and arginine decarboxylase (*SP_0916*), which produces agmatine and is required for capsule production (43). Interestingly, the pilus locus that encode proteins for synthesis of a multi subunit surface structure involved in interaction with host cellular components (39) was upregulated during co-infection (**Fig. 4B, Table S1 and S2**), suggesting that pro-inflammatory conditions in the URT potentially stimulate the expression of the type 1 pilus. We had previously demonstrated that the pneumococcal pilus present in a subset of isolates does not impact *Spn* shedding and colonization during mono-infection (24). To determine if it is important during co-infection, we tested wild-type and an isogenic pilus deficient mutant (*Δpilus*) during mono- and IAV co-infection (**Fig. 5**). In concordance with our earlier results, we did not observe any shedding differences in either daily or pooled (combined 5 days of shedding values and plotted them together) during mono-infection (**Fig. 5A**). During IAV co-infection, the daily shedding data demonstrated that both strains shed equally well over the course of the study (**Fig. 5Bi**). However, when the shedding data are combined, we observed that high shedding events (> 750) occurred less frequently from pups inoculated with the *Δpilus* mutant (**Fig. 5Bii)**. Both the WT and *Δpilus* strains colonized equally well under both mono- and co-infection conditions (**Fig S5**), suggesting that the reduction in high shedding events observed with IAV co-infection are not due to a defect in ability of the strains to colonize the nasopharynx.

**Figure 4.**
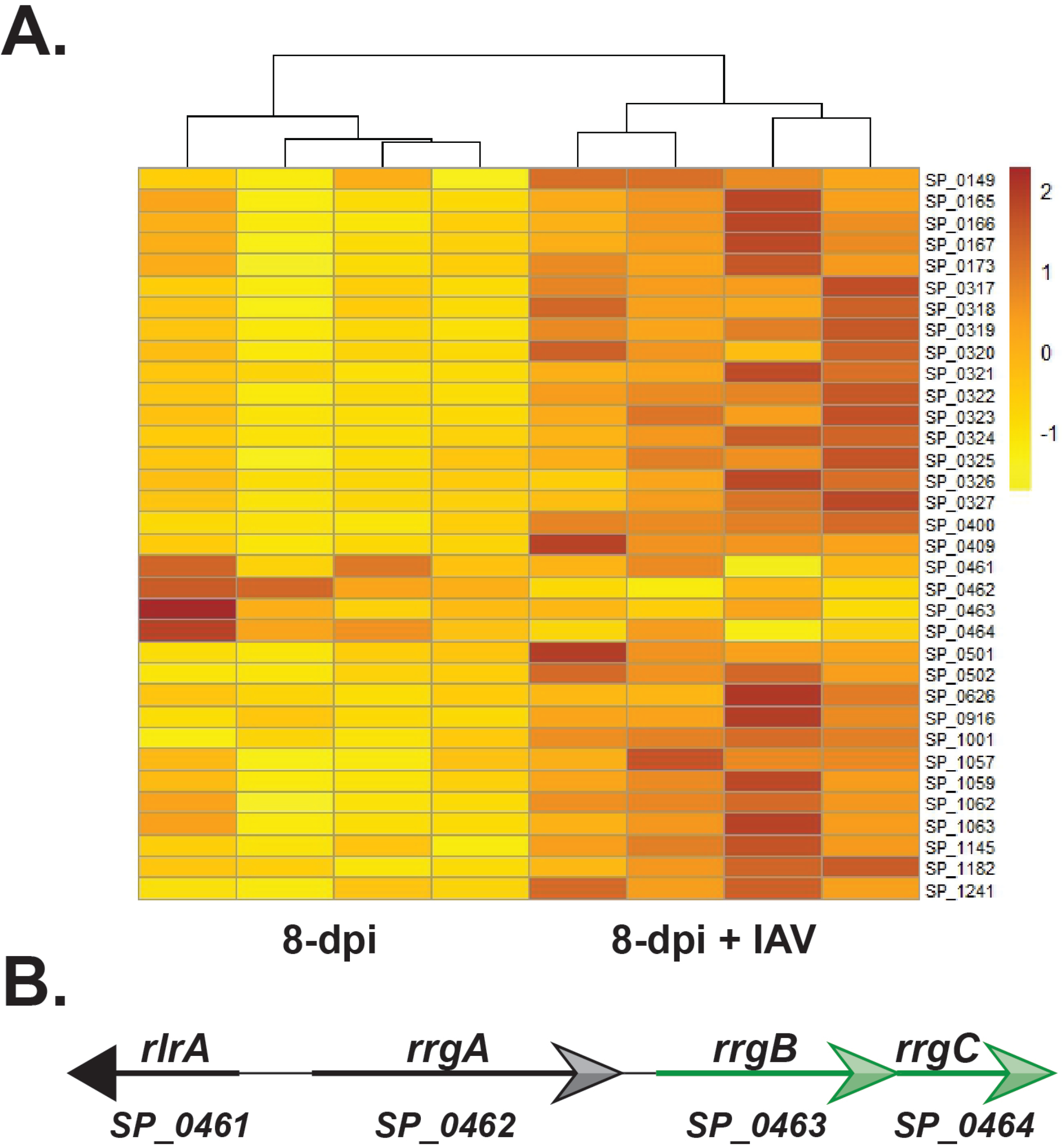
Heatmap of differentially expressed *S. pneumoniae* genes during mono and IAV coinfection. **(A)** Shown are top 30 genes that are above the *P*_*adj*_ cutoff value that are upregulated during *S. pneumoniae*-IAV coinfection in comparison to *S. pneumoniae* coinfection. Besides the 30 genes included are the genes that include the type 1 pilus islet. Expression values were extracted directly from the deseq2 analysis and represent raw values. The scale bar adjacent to the heatmap indicates the numeric range of these values, facilitating direct comparison of expression levels between samples. **(B)** Schematic representation of the *S. pneumoniae* TIGR4 pilus island 1. Locus tag and gene name are included. Genes highlighted in green are those that were upregulated during coinfection.

**Fig 5.**
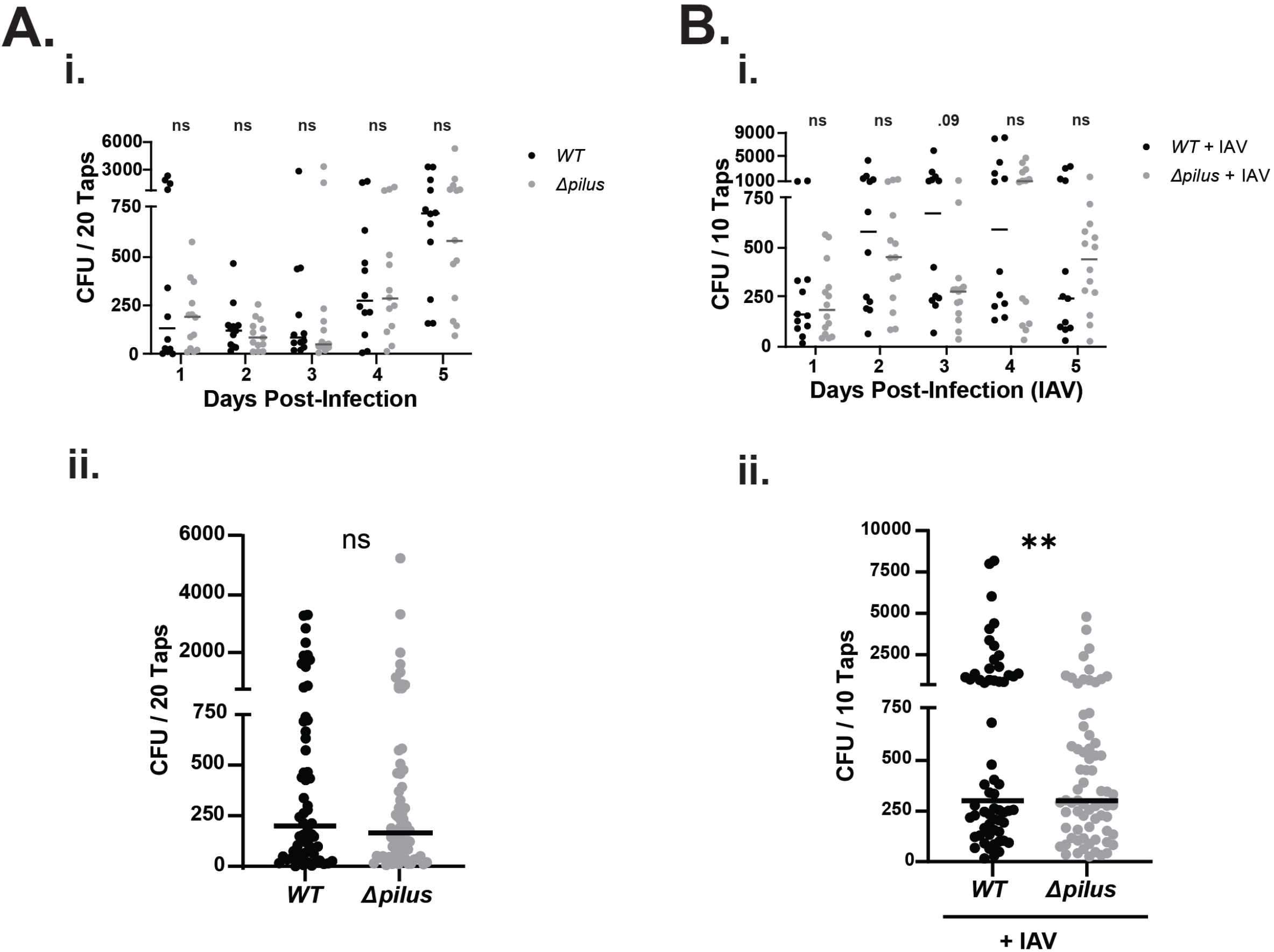
The Pilus contributes towards *S. pneumoniae* high shedding events during a concurrent influenza A (IAV) infection. **(A) (i.)** 4 day old pups were challenged i.n with the indicated strain, and nasal secretion were collected and bacterial burden determined for 5 days P.I. Median CFU values for each day are indicated, each symbol represents the number of CFUs measured from a single pup (n ≥ 5 for each strain tested). **(ii)** Pooled 5 days of median shedding for the two strains is shown. Shedding differences between the two strains including high shedding events (> 750) were calculated using the Mann-Whitney *U* test. **(B)** Pups were challenged i.n at 4 of age with the indicated construct and challenged with IAV at day 9 of life and bacterial shedding from nasal secretions were quantified for 5 days post IAV infection, with median CFU values shown for each day (n ≥ 5 for each strain tested). **(ii)** Pooled 5 days of median shedding for the two strains is shown. Shedding differences between the two strains including high shedding events (> 750) were calculated using the Mann-Whitney *U* test. Median values are shown, with statistical differences calculated using Mann Whitney *U* test. *\*\*P < 0*.*01;* ns, not significant.

### The *Spn* pilus locus is a pro-inflammatory determinant during coinfection

Inflammation has been demonstrated to be a key determinant for robust pneumococcal shedding from the nasopharynx (12). Therefore, we examined whether the reduced shedding of the *Δpilus* mutant from the URT was due to reduced inflammation during coinfection of IAV and Δ*pilus* compared to IAV and wild-type. Inflammatory markers such as chemokines (CXCL-1 and 2) and cytokines (IL-1β) are considered hallmarks of for *Spn*-induced inflammation in the URT. Thus, we measured their transcript levels in the nasopharynx on Day 8 P.I with a concurrent IAV infection. We observed reduced transcript levels of CXCL-2 and IL-1β (**Fig 6A**) from mice inoculated with Δ*pilus*, suggesting that differences in URT inflammation likely account for the variation in number of high shedding events.

**Figure 6.**
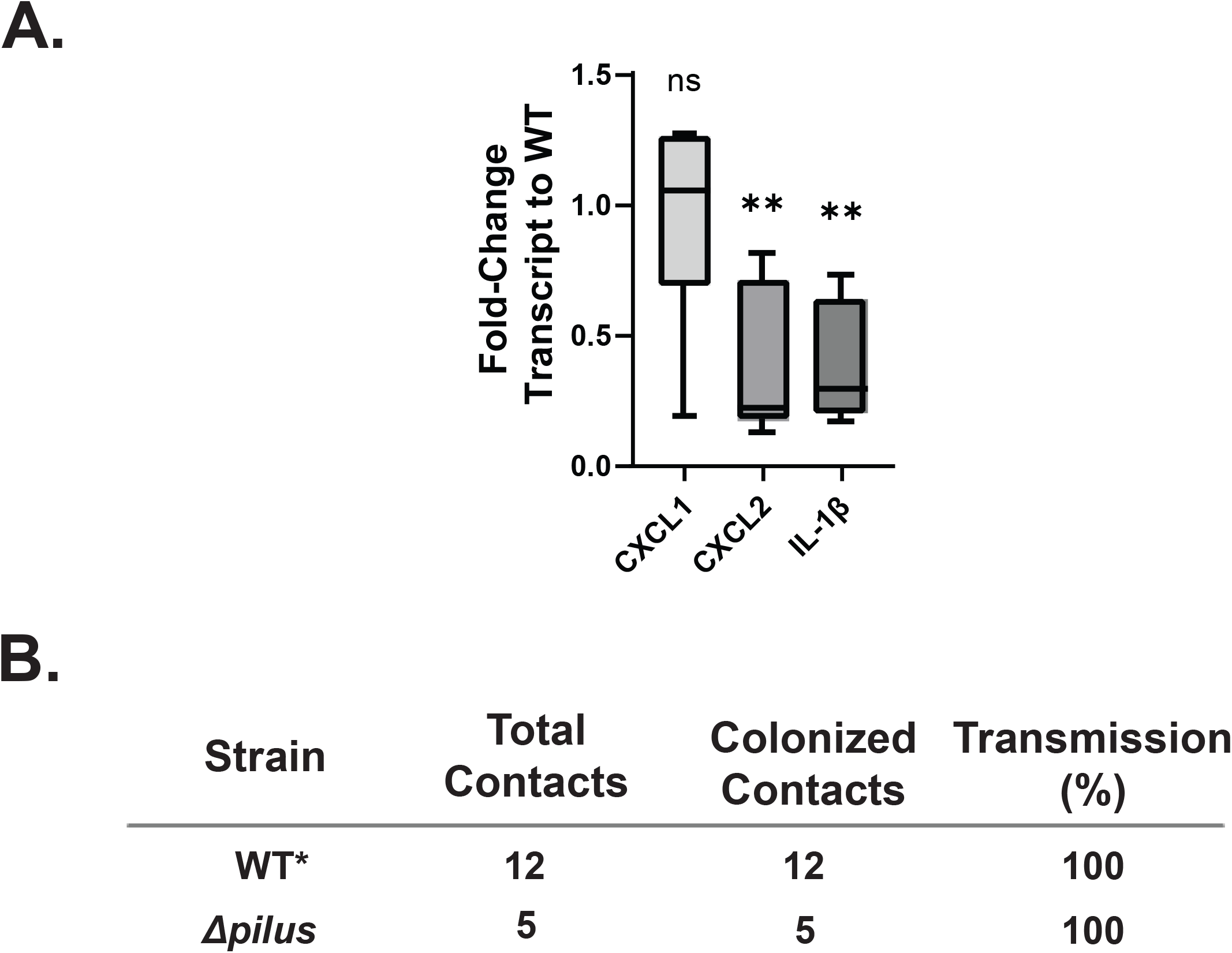
The absence of Pilus locus leads to reduced URT inflammation during a concurrent *S. pneumoniae*-IAV infection. **(A)** Quantitative real-time PCR was used to examine host gene transcripts from pups colonized with the pilus-deletion mutant (*Δpilus*) relative to that in the WT strain-inoculated pups with a concurrent IAV infection at age 12 days. Values are ±SEM (n>5 pups). Statistical differences between the WT and *Δpilus* strain were calculated using the Mann-Whitney *U* test. **(B)** Summary of the rate of transmission of the *Δpilus* strain and the results from the WT (*historic data) (31) from pups that were colonized index pups with *S. pneumoniae* at age of 4 days (at a 1:4 ratio of index to contact mice). 5 days later all the pups in the litter received IAV, and the transmission efficiency determined at age of 14 days. Fisher’s exact text was used for statistical comparison. *\*\*P < 0*.*01;* ns, not significant.

Next, as high shedding is considered a pre-requisite for transmission (12, 30), and as IAV coinfection causes higher transmission events compared to *Spn* mono-infection (31), we determined whether the reduction in high shedding events from mice inoculated with the Δ*pilus* mutant correlate with reduced transmission rates during IAV coinfection. One in three mice per litter was designated an index pup and colonized with either the wild-type or mutant strain at day 4 of life. On day 9 of life all the pups in the litter were infected with IAV. The pups were housed together with the dam for the entire duration of the transmission study. At day 14 of life all the pups were euthanized, and retrograde lavages were cultured to determine whether any transmission events had occurred between the index and the contact pups. However, transmission frequency of the *Δpilus* mutant was similar to that previously reported for the wild-type strain(31) (**Fig 6B**). Thus, while the Δ*pilus* mutant had fewer high shedding events, the reduced shedding did not correlate with reduced transmission events.

## Discussion

Here, we leveraged our murine model of *Spn* colonization, shedding, and host-to-host transmission to provide insight into the global transcriptional changes that occur during *Spn* colonization of the upper respiratory tract. Our data show that the transcriptional profiles of bacteria isolated from day 1 P.I and day 8 P.I are generally similar; however, these profiles differ considerably when IAV is introduced as a coinfecting agent. Many of the genes identified in these three conditions match those from an earlier transposon mutagenesis screen in this same murine model that identified genes important for *Spn* shedding and colonization(30, 34).

Furthermore, we demonstrate that the pilus locus of the isolate TIGR4 is dispensable for colonization during both *Spn* and IAV-*Spn* infections. However, while no overall differences in median shedding were observed under mono-infection and co-infection, we did observe that the high shedding events during co-infection were significantly reduced in the pilus mutant, which correlated with reduced inflammation as measured by transcript levels of chemokines and cytokines in the upper respiratory tract. Lastly, we did not observe corresponding differences in transmission efficiency from the reduced shedding. However, our transmission model assumes transmission through interactions with nasal secretions rather than aerosol (12), and may lack the sensitivity to detect the correlation between high shedding events and host-to-host transmission.

Previous RNA-seq approaches have primarily focused on using *in vitro* conditions that mimic the *in vivo* environments that *Spn* encounters in the host such as the lung, cerebrospinal fluid, blood, and mucin(7, 41, 44-47). Other high-throughput RNA-seq studies have focused on both bacterial and host responses, either *in vitro* or *in vivo*(41, 45, 46, 48). More recently, seminal work by D’Mello et al.,(28) aimed to provide a molecular understanding of the transcriptional changes *Spn* undergoes at different anatomical sites in adult mice, results that align with our study using infant mice. By comparing two different *Spn* mono-infection timepoints, as well as mono-versus IAV co-infection, our study provides novel insight into transcriptional modulations that have not previously been examined. Many of the pneumococcal genes identified in these screens, including glycan cleavage, uptake, metabolism, bacteriocins, and other metabolic genes, were also identified in our screen, providing robustness to our results.

A previous transposon mutagenesis screen in the infant mouse model identified both shedding (transmission) and colonization determinants for *Spn* (30, 34). In our current transcriptomics approach we identified 17 genes that were either required for pneumococcal shedding or for robust colonization (**Fig 2**). Three loci (*SP_0060-0064*; *SP_0317-320*; *SP_0321-0325*), were identified as significantly upregulated for both day 1 and day 8 P.I. *SP_0060-0064* encodes a surface beta-galactosidase (*bgaC*, involved in glycan degradation and required for nasopharyngeal colonization (49) and the mannose-type PTS system, and genes in this locus were upregulated when grown in mucin-containing media. Interestingly, *SP_0317-320* and *SP_0321-0325* are present only in pathogenic *Streptococcus* species, suggesting their importance for pneumococcal pathogenesis (50). These loci contain genes that encode enzymes that, together with hyaluronate lyase (*SP_0314*; *hyl*), are involved in the uptake and metabolism of hyaluronic acid, which is present on the apical surface of epithelial cells and a component of the capsule for some bacterial species. *Spn* is known to use hyaluronic acid as a carbon source, including from bacteria whose capsules contain hyaluronic acid. Inability to metabolize hyaluronic acid is associated with poor URT colonization, underscoring the importance of this system to *Spn* colonization (32, 42, 51). Additionally, *SP_2075* (*patA*) was highly upregulated during coinfection and previously identified as a colonization determinant (34). *patA* and *patB* encode the PatAB multidrug efflux system that has been demonstrated to be important for resistance against fluoroquinolones (52), and it is highly expressed in clinical isolates (53). However, further studies are required to determine the physiological importance of this system during *Spn* colonization of the URT.

Because of its high expression levels during co-infection, we focused on the pneumococcal Pilus islet-1. This structure is only present in a subset of *Spn* isolates, including TIGR4, and is a multi-subunit structure that contributes to invasive disease (39). RNA-seq from mono-infected mice demonstrated that in comparison to *in vitro* conditions, the Pilus locus expression was downregulated, consistent with previous reports showing that Pilus is dispensable for pneumococcal shedding and colonization in infant and adult murine models (24, 54). However, the role for Pilus during IAV coinfection had not been explored. While the Δ*pilus* mutant had no defect in colonization, the number of high shedding events was significantly reduced in the Δ*pilus* mutant, suggesting that it likely contributes to *Spn* shedding during IAV coinfection. Previous work has demonstrated that TLR2 is known to recognize cellular components of pneumococcus and contributes to the inflammatory response that drives *Spn* shedding (30) and it has been established that the oligomerized form of RrgA, the tip adhesin, is a Toll-like receptor (TLR) 2 agonist (55). Given that inflammation plays a leading role in pneumococcal shedding and the Pilus induces the host inflammatory response (54), we then examined whether the low shedding events during *Spn*-IAV coinfection was due to changes in inflammation. The reduced levels of chemokine and cytokine transcripts in mice inoculated with Δ*pilus* suggests that the Pilus contributes to increased inflammation in the URT during a concurrent IAV infection. Thus, we postulate that the Pilus increases inflammation via TLR2 recognition, and the high inflammation in response to both IAV and the Pilus results in an increased number of high-shedding events. Recent epidemiological data suggest that the Pilus islet-1 strains are also strongly associated with high levels of antibiotic resistance (56). Thus, it would be interesting to determine epidemiologically whether Pilus islet-1 strains with concurrent IAV infection exhibit a higher transmission frequency in a community setting.

One caveat of our study was that the infection model uses infant mice, which are not considered a natural reservoir for *Spn*. Further validation of the genes identified in the screen in either the human challenge or the ferret model of *Spn* infection (57) would provide stronger physiologically relevant insight. Only a single isolate of *Spn*, the invasive strain TIGR4, was used in these studies, as it is well annotated and has been widely used in the research community (26). Other strains associated with carriage that do not include the pilus-1 locus may have a different transcriptional profile in the nasopharynx. Lastly, we used rich broth media as a control in our studies. Our results could differ depending on the control media used. However, we identified many genes required for glycan uptake and metabolism, as well as other metabolic genes previously ascertained to be important in nasopharyngeal colonization, and we also compared transcriptional profiles between mono-infection and co-infection, thereby strengthening our study.

Although not without caveats, our study provides key insight into the transcriptional modulation of *Spn* during colonization of the upper respiratory tract and furthers our understanding of IAV-mediated *Spn* transcriptional changes. Our transcriptomics results will serve as a valuable resource for both the pneumococcal and the broader pathogenesis communities. This work also provides foundational data for identifying potential novel therapeutic targets that are much needed to reduce the health care burden of *Spn*.

## Materials and Methods

### Ethics statement

This study was conducted in accordance with the animal welfare guidelines established by the National Science Foundation and the Public Health Service Policy on Humane Care and Use of Laboratory Animals. All procedures involving animals were carried out following the standards set by the American Association for Laboratory Animal Science (AALAS) and were approved by the Institutional Animal Care and Use Committee (IACUC) at Wake Forest Baptist Medical Center. The approved protocol number for this project is A23-039.

### Growth conditions and strain construction

Strains used in this study include a streptomycin derivative of TIGR4 (serotype 4 isolate) (24) and an isogenic pilus islet locus mutant (Δ*rlrA;* ΔPilus) (24). Pneumococcal Strains were grown in a 37°C circulating water bath in tryptic soy broth (Becton, Dickinson [BD]) supplemented with 0.3% yeast extract (TSY) until optical density at 620nm (OD_620_) of 0.4. Once *Spn* cultures reached desired OD_620_ they were diluted in sterile phosphate-buffer saline (PBS) for intranasal inoculation and other assays. To quantitate *S. pneumoniae*, serial dilutions were plated on TSY agar-streptomycin (200 μg/mL), supplemented with either catalase (6,300 U/plate; Worthington Biochemical Corporation) or 5% defibrinated sheep’s blood, and incubated overnight at 37°C with 5% CO_2_.

### Infant mouse infection model for shedding, colonization, and transmission

All mouse experiments were conducted using specific pathogen-free (SPF) C57BL/6J mice, originally obtained from The Jackson Laboratory (Bar Harbor, ME). These mice were subsequently bred and maintained in the animal facility at Biotech Place, Wake Forest Baptist Medical Center. Throughout the study, pups remained housed with their dam (mother) and exhibited normal weight gain comparable to uninfected controls.

Mouse infections were done on day 4 of life. Pups were intranasally infected with ∼2000 CFU of *Spn* in 3μL of PBS. To enumerate nasal shedding of *Spn* post-inoculation, the nares of the infected pups were gently tapped (20 taps per pup for *Spn* mono-infection, 10 taps per pups exposed to IAV secondary-infection) onto a TSY agar plate supplemented with streptomycin (200 μg/mL) and catalase to only allow for the growth of *Spn*. The nasal secretions were spread using a sterile cotton swab, and the plates were incubated as previously described (15, 24). Due to natural day-to-day variability in shedding, CFU values from individual pups were pooled across the 5-day study period. The IAV virus used in co-infection studies is the mouse-adapted influenza A virus/Puerto Rico/8/34(H1N1), here referred to as “PR8”. Five days after primary *Spn* challenge (day 9 of life), intranasal IAV secondary challenge was given with 2 × 10^4^ PFU of PR8 suspended in 3µL of PBS. Daily shedding was quantified as described above.

At the conclusion of the study, pups were euthanized at designated time points using CO2 asphyxiation followed by cardiac puncture. The upper respiratory tract (URT) was then exposed and lavaged with 200 μL of sterile PBS, introduced via a needle inserted into the trachea, with the lavage fluid collected from the nares. The resulting nasal wash was serially diluted and plated on TSY agar supplemented with streptomycin (200 μg/mL) and catalase to assess colonization density. The detection limit for *Spn* in the lavage fluid was 33 CFU/mL.

For transmission studies the *S. pneumoniae*-IAV coinfection transmission model was followed as described previously (24, 31). Briefly, one in three pups in a litter was randomly selected at day 4 of life and inoculated with the Δ*rlrA* mutant. These index pups were then returned to their litter mates (contact) and dam. At day 9 of life, all the pups in the litter were inoculated with IAV/HKx31 as mentioned above. To determine *S. pneumoniae* transmission frequency, the index and contact pups were euthanized at day 14 of life, and colonization density assessed by sampling nasal lavages.

### *In vivo* and *in vitro* RNA isolation

Bacterial RNA was isolated from the upper respiratory tract from *S. pneumoniae* (TIGR4) colonized pups using a modified acid phenol chloroform method (58). Briefly, retrograde nasal lavages in PBS were collected from colonized pups at designated time points and immediately incubated with the RNAprotect (Qiagen) and processed according to manufacturer’s instructions. Processed lavage samples from two pups were pooled together and then resuspended in prewarmed (65°C) phenol/chloroform/isoamyl alcohol (pH 4.5; Ambion 9722) and incubated for 5 minutes at 65°C. To this prewarmed 300 µl of NAES (50 mM sodium acetate pH 5.1, 10 mM EDTA, 1% SDS) was added, and sample further incubated at 65°C with intermittent mixing. The mixture was cooled on ice for 1 minute and the phases separated by centrifugation at 15,500 *x g* for 5 minutes at room temperature. Potassium acetate (pH 5.5) was added to the aqueous phase giving a final concentration of 300 mM, and RNA precipitated overnight using 2.5 volume ice cold ethanol with 40 ng glycogen µl^-1^ (Sigma 1767). The precipitated RNA was pelleted via centrifugation at 15,500 *x g* for 10 minutes at 4°C. The pellet was further washed with 70% ethanol, air dried and resuspended in nuclease free water. The samples were further purified using the RNeasy kit (Qiagen) per the manufacturer’s instructions. RNA from mid-log phase (OD_620_ = 0.4) *S. pneumoniae* grown statically in TSY medium were processed similarly as described above.

### cDNA Library Preparation, sequencing and RNA-seq analyses

Total RNA was used to construct stranded cDNA libraries using the Illumina® Stranded Total RNA Prep, Ligation with Ribo-Zero Plus Microbiome kit. Ribosomal RNA was first depleted, followed by enzymatic fragmentation, reverse transcription, and purification of double-stranded cDNA using AMPure XP magnetic beads. The cDNA fragments were end-repaired, 3′ adenylated, and ligated with Illumina sequencing adapters, followed by PCR pre-amplification to generate stranded libraries. Library size distribution and quality were assessed on the Agilent 4200 TapeStation, and cDNA concentrations were quantified using the Qubit 3.0 Fluorometer (Thermo Fisher Scientific, USA). Final libraries were pooled and sequenced on the Illumina NovaSeq 6000 using the NovaSeq 6000 SP Reagent Kit v1.5 (100 cycles) flow cell.

All samples achieved a sequencing depth exceeding 12 million reads and were evaluated for quality by FastQC (Babraham Bioinformatics). Reads were aligned to the *Streptococcus pneumoniae* TIGR4 genome assembly (ASM688v1) via the STAR sequence aligner (59) with unspliced alignment function and summarized by featureCounts(60). Differential expression analysis was conducted using DESeq2(61). Genes with log2 fold change ≥1 (twofold change) and adjusted *P* value for multiple comparisons of ≤0.05 were considered as being differentially expressed in our analysis. Pneumococcal gene pathways used in our analysis were taken from the Kyoto Encyclopedia of Genes and Genomes (KEGG).

### Host RNA isolation and qRT-PCR

Host RNA was isolated by lavaging the upper respiratory tract (URT) of pups at designated time points that were initially inoculated with *S. pneumoniae* (day 4 of life) and subsequently inoculated with IAV or PBS (mock) at day 9 of life, using RLT lysis buffer (Qiagen) to isolate host RNA. RNA purification (Qiagen) and cDNA synthesis (Bio-Rad) were performed according to the manufacturer’s instructions. Quantitative real-time PCR (qRT-PCR) was carried out using Bio-Rad SYBR Green Master Mix, with 2.5 ng of cDNA and 0.5 μM primers per reaction. Primers targeting glyceraldehyde-3-phosphate dehydrogenase (GAPDH) were used as an internal control. Reactions were run in duplicate on a CFX384 Touch Real-Time PCR Detection System (Bio-Rad), and each experiment was repeated twice. Gene expression levels were quantified using the *ΔΔC*_*T*_ threshold cycle (C_T_). Primer sequences used in this study were previously described(62, 63).

### Statistical analysis

The ggplot2 package in R version 4.4.2 was used to conduct principal component analysis as well as create heatmaps and figures for pathway analysis.

Statistical analyses were performed using GraphPad Prism (version 10.0) software (GraphPad Software, Inc., San Diego, CA).

## Data availability

The raw FASTQ reads have been submitted to the NCBI’s Sequence Read Archive (SRA) under the BioProject PRJNA1464539 BioSample ID -SAMN59720126-SAMN59720141 with GEO accession GSE330514.

## Acknowledgements

We are grateful to Jeffrey N. Weiser (NYU School of Medicine) for providing us with the strains and in performing the transmission studies. We further thank Dr. Kimberly A. Walker (UNC Chapel Hill) for providing critical feedback on this manuscript. This project was supported by startup funds provided by Wake Forest Baptist Medical Center and U.S. Public Health Service grant (AI154047) to M.A.Z.

We also acknowledge the support of the Wake Forest Baptist Comprehensive Cancer Center Genomics Shared Resources, supported by the National Cancer Institute’s Cancer Support Grant (P30CA012197).

---

**Supplemental Figure 1. (A)** *Spn* colonization density with and without a concurrent IAV infection obtained in URT lavage fluids obtained from pups at either day 9 of age (monoinfection) or at the indicated timepoint post IAV infection (coinfection). **(B-D)** Box and whiskers graph depicting quantitative real-time PCR results for host gene transcripts from pups colonized with the WT strain (monoinfection) relative to that in the WT strain-inoculated pups with a concurrent IAV infection at either 24 hours **(B)**, 72 hours **(C)** or 120 hours **(D)**. (n≥3 pups).

**Supplemental Figure 2**. RNA read counts of TIGR4Sm^R^ across the three time-points and broth and *in vivo* samples. Also shown is the alignment summary of both *in vivo* and *in vitro* samples.

**Supplemental Figure 3**. Heatmaps depicting the top 30 pneumococcal genes with locus tags included that are downregulated when comparing **(A)** day 1 P.I and TSY, **(B)** day 8 P.I and TSY, and **(C)** day 8 P.I with a concurrent IAV infection and TSY.

**Supplemental Figure 4**. Venn diagram displaying the number of genes in a particular KEGG global pathway, with upregulated genes in parenthesis, with overlap shown for genes involved in multiple KEGG pathways, comparing **(A)** day 1 P.I to broth, **(B)** day 8 P.I to broth and, **(C)** day 8 P.I with a concurrent IAV infection.

**Supplemental Figure 5**. Colonization density of the indicated strains in cultures obtained in URT lavage fluids obtained from pups at either day 9 of age (monoinfection) or day 14 of age (coinfection). Median values are shown, with statistical differences calculated using Mann Whitney *U* test. n≥5 pups. ns, not significant.

**Supplemental Table 1**. Pneumococcal gene expression during upper respiratory tract colonization.

**Supplemental Table 2**. Top 30 pneumococcal genes with locus tags included that are either upregulated or downregulated.

